# Differential effects of high versus low linear energy transfer (LET) radiation on type-I interferon (IFNβ) and TREX1 responses

**DOI:** 10.1101/2021.07.07.451516

**Authors:** Devin Miles, Ning Cao, George Sandison, Robert D. Stewart, Greg Moffitt, Thomas Pulliam, Upendra Parvathaneni, Peter Goff, Paul Nghiem, Keith Stantz

## Abstract

**Purpose:** Cancer cells produce innate immune signals following radiation damage, with STING pathway signaling as a critical mediator. High linear energy transfer (LET) radiations create larger numbers of DNA double-strand breaks (DSBs) per unit dose than low-LET radiations and may therefore be more immunogenic. We studied the dose response characteristics of pro-immunogenic type-I interferon, interferon-beta (IFNβ), and its reported suppressor signal, three-prime repair exonuclease 1 (TREX1), *in vitro* with low-LET x-rays and high-LET fast neutrons.

**Methods:** Merkel cell carcinoma cells (MCC) were irradiated by graded doses of x-rays (1-24 Gy) or fast neutrons (1-8 Gy). IFNβ was measured as a function of dose via ELISA assay, and exonuclease TREX1 expression via immunofluorescence microscopy. The Monte Carlo damage simulation (MCDS) was used to model fast neutron relative biological effectiveness for DSB induction (RBE_DSB_) and compared to laboratory measurements of the RBE for IFNβ production (RBE_IFNβ_) and TREX1 upregulation (RBE_TREX1_). RBE_IFNβ_ models were also applied to radiation transport simulations to quantify the potential secretion of IFNβ in representative clinical beams.

**Results:** Peak IFNβ secretion occurred at 5.7 Gy for fast neutrons and at 14.0 Gy for x-rays, i.e., an effective RBE_IFNβ_ of 2.5 ± 0.2. The amplitude (peak value) of secreted IFNβ signal did not significantly differ between x-rays and fast neutrons (P > 0.05). TREX1 signal increased linearly with absorbed dose, with a four-fold higher upregulation per unit dose for fast neutrons relative to x-rays (RBE_TREX1_ of 4.0 ± 0.1). Monte Carlo modeling of IFNβ suggests Bragg peak-to-entrance ratios of IFNβ production of 40, 100, and 120 for proton, alpha, and carbon ion beams, respectively, a factor of 10-20-fold higher compared to their corresponding physical dose peak-to-entrance ratios. The spatial width of the Bragg peak for IFNβ production is also a factor of two smaller.

**Conclusion:** High-LET fast neutrons initiate a larger IFNβ response per unit absorbed dose than low-LET x-rays (i.e., RBE_IFNβ_ value of 2.5). The RBE value for IFNβ is quite similar to data reported in the literature for DSB induction and cellular, post-irradiation micronucleation formation for neutrons and x-rays. The increased IFNβ release after high-LET radiation may be a contributing factor in stimulating a systemic anti-tumor, adaptive immune response (abscopal effect). However, our results indicate that TREX1 anti-inflammatory signaling *in vitro* for MCC cells is larger per unit dose for fast neutrons than for x-rays (RBE_TREX1_ of 4.0). Given these competing effects, additional studies are needed to clarify whether or not high-LET radiations are therapeutically advantageous over low-LET radiation for pro-inflammatory immune signaling in other cell lines *in vitro* and for *in vivo* cancer models.

## Introduction

Recent radiotherapy research has demonstrated an increase in anti-tumor immune activity following hypofractionated radiation dose delivery using low-LET radiation sources^1,2^. Cellular damage by radiation may result in accumulation of genomic DNA in the cytosol prompting a subsequent cascade of interactions resulting in the secretion of type-I interferons (IFNs), such as IFNβ. IFNβ promotes a localized immune response leading to an increased prevalence of tumor-infiltrating lymphocytes. These immune cells, in turn, can initiate anti-cancer immunity through the release of the type-II IFN, IFNγ, a cytokine catalyzing antigen presentation to T-cells by upregulating MHC I in target cancer cells and MHC II on antigen presenting cells (APCs).^3^ Initiation of these former cellular responses (IFNβ) from cancer cells can in part be attributed to activation of the innate cellular response mechanism known as the Stimulator of Interferon Genes (STING) signaling pathway, which is most often triggered in humans following cellular recognition of DNA in cytosol following an intracellular viral infection.^4,5^ Thus, radiation-induced DNA damage may lead to signaling that engages the adaptive immune system.

A radiation-dose-dependent induced immunogenic response has been observed in human and murine cancer cell lines. Such a response may be optimal when cells are sufficiently damaged to create enough cytosolic DNA to activate STING and secrete IFNβ, but not too much DNA damage to suppress STING activation by triggering the upregulation of TREX1.^6,7^ *In vivo*, a low-LET x-ray dose of about 8 Gy, when delivered in combination with immune checkpoint inhibitors (ICI), appears optimal for inducing a systemic immune response in murine models of melanoma, colon carcinoma, and adenocarcinoma.^1,6,7,8^ Immunogenic responses at similar doses have also been observed in patients using megavoltage (MV) x-rays and have been shown to improve patient outcome.^9,10^ Alternatively, a number of preclinical and clinical studies have indicated that high-LET radiation is highly immunogenic at lower doses, including studies with alpha particles,^11^ carbon ions,^12,13^ and fast neutrons.^14^ For example, a recent case study observed an out-of-field (abscopal) response in a Merkel cell carcinoma (MCC) patient, previously refractory to low-LET radiation and PD-1 ICI with pembrolizumab, following high-LET fast neutron therapy (2 daily fractions of 3 Gy).^14^ This case study highlights the potential of high-LET radiation to overcome metastatic disease that is non-responsive to low-LET radiation or immunotherapy alone. These studies provide tantalizing, although anecdotal, evidence suggesting that high-LET radiation therapy, with doses low enough to be easily tolerated by most normal tissues, is potentially more effective than low-LET radiation at initiating an out-of-field, tumor-specific immune response. To date, the dose response of IFNβ for any high-LET radiation has not been reported in the literature.

In this study, we investigate the *in vitro* dose-response of IFNβ and TREX1 in MCC following irradiation by low-LET x-rays and high-LET fast neutrons, from which an RBE (relative biological effectiveness) with respect to IFNβ secretion (RBE_IFNβ_) and TREX1 upregulation (RBE_TREX1_) was defined. We also used Monte Carlo modeling of DNA damage to develop mechanism-inspired empirical models of IFNβ and TREX1 regulation as a function of absorbed dose and the RBE for DSB induction (RBE_DSB_). These models were applied to general particle transport simulation to investigate the potential secretion of IFNβ for clinically relevant hadron therapy beams. The data from these experiments and Monte Carlo simulations are a useful guide for additional studies of pro-immunogenic radiotherapy treatments and future clinical trials.

## Materials and Methods

### In vitro experimental design

Merkel cell carcinoma (MCC13) cells (a Merkel polyomavirus-negative cell line)^17,18^ were cultured in RPMI 1640 with L-glutamine, supplemented with 10% heat-inactivated fetal bovine serum and penicillin/streptomycin. Short tandem repeat analyses were used to verify the authenticity of this cell line prior to experimentation. Confirmation of intact cGAS-STING was performed by IFNβ measurement following transfection of non-sheared calf thymus DNA using Lipofectamine 2000. These cells tested negative for mycoplasma contamination.

To test the endpoint of IFNβ production, 6×10^5^ MCC13 cells were seeded onto 6 cm dishes with a total of 3.8 mL growth medium one day prior to irradiation. Cells were irradiated at room temperature using 220 kVp x-rays (0.8 mm Be and 0.15 mm Cu filtration) on an Xstrahl SARRP (Xstrahl Inc, Suwanee GA) or using 50.5 MV fast neutrons from the Clinical Neutron Therapy System (CNTS) at the University of Washington,^15,16^ with delivered doses escalating up to 24 Gy (SARRP) or up to 8 Gy (CNTS). Growth medium was replaced immediately following irradiation, and cells were allowed to incubate at 37°C for 96 hours. The incubation time based on the MCC13 doubling time *in vitro*, where the exposed cells are expected to undergo at least two cell cycles, allowing for micronucleus formation and collapse. Sampling at earlier time points resulted in reduced measurement amplitudes. Following incubation, cell supernatant was harvested and centrifuged to remove dead cells and debris. Cell-free supernatant was aliquoted and stored at −20°C until assay. IFNβ concentrations were measured using Duoset human IFNβ ELISA kit with ancillary supplies (R&D Systems, DY814-05) following the product protocol. Optical densities were measured using a Synergy4 microplate reader immediately after assay completion and analyzed in MATLAB using in-house developed scripts. IFNβ quantification was performed using an IFNβ standard curve, using a 7-point twofold dilution of recombinant human IFNβ in RPMI, and readout was normalized to the number of cells per plate.

For exonuclease TREX1 assay, cells were seeded into 4-well chamber slides (Thermo Fisher Scientific, Nunc Lab-Tek II) one day prior to irradiation at a density of 4.5×10^4^ cells/well, with a total of 0.4mL of medium. Cells were irradiated as described above and slides were allowed 96 hours to incubate at 37°C. Following incubation, cells were fixed with 4% paraformaldehyde, permeabilized with 0.5% Tween-20, and blocked with 3% bovine serum albumin. Immunofluorescence staining was performed using a pre-conjugated Alexa594/TREX1 antibody (Abcam ab217095) diluted 1:500, with a 2-hour incubation time at room temperature. Coverslips were mounted using ProGold with DAPI counterstain. Immunofluorescence imaging was performed using a Nikon 90i microscope at 20X magnification using Texas Red filtration. The average integrated density (product of average intensity and cell area) per cell was quantified using ImageJ and analyzed in MATLAB.

The above experiments were repeated three times (n=3) and the mean and standard deviation was determined for each dose. For all experiments, a paired t-test was used to determine statistical significance.

### Modeling of DSB Induction and IFNβ and TREX1 Response

To develop a dose-response model of *in vitro* STING regulation and suppression, IFNβ and TREX1 dose response curves from x-ray and neutron irradiations were fitted using MATLAB to a non-linear function (Eqn. 1) or a first-order polynomial (Eqn. 2), respectively.

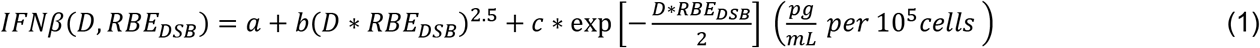

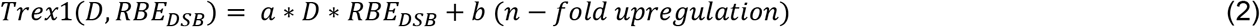

The measured data were independently fit to Eqn (1) and (2) as functions of absorbed dose and a Monte Carlo calculated RBE for DSB induction (RBE_DSB_) relative to Colbalt-60 γ-rays.^19–21^ RBE_DSB_ was determined using the Monte Carlo Damage Simulation (MCDS), which develops nucleotide-level maps of the clusters of DNA lesions formed by low and high-LET charged particles and the clustering of DNA lesions into DSB and other types of clusters (i.e., clusters of base damage and single-strand breaks). DNA damage for x-rays and fast neutrons were simulated by combining information from the MCDS with a larger-scale Monte Carlo radiation transport.^22^ All MCDS simulations were performed using 50,000 trials to achieve a standard error of less than 1%, using default settings for nucleus diameter (5 μm) and DNA content (1 Gbp).

Monte Carlo simulations by our group indicate an RBE_DSB_ for SARRP x-rays of 1.17. This RBE was computed using the MCDS software with the secondary electron energy fluence from a 220 kV x-ray source, as input, calculated using the FLUKA general purpose Monte Carlo software. Briefly, our 220 kV photon source was modeled to match manufacturer specifications for a Varian NDI 225/22 x-ray tube, with 0.8 mm beryllium and 0.15 mm copper filtration. PRECISIO default parameters were implemented, with photon and electron production and transport cutoffs set to 1 keV. Comparable Monte Carlo simulations using MCNP and MCDS give a very similar RBE_DSB_ of 1.20 for x-rays produced within SARRP. Monte Carlo simulations of CNTS fast neutrons relative to ^60^Co γ-rays suggest an RBE_DSB_ in the range from 2.5 to 3.0.^21^ For CNTS neutrons relative to SARRP x-rays, the estimated RBE_DSB_ is 2.09 to 2.50.

### Simulating IFNβ and TREX1 production for various clinical radiation beams

We also performed Monte Carlo simulations for other clinically relevant radiation sources, including 6 MV x-rays and charged particle beams (proton, alpha, and carbon ions), to quantify the RBE for the endpoints of: DSB induction (RBE_DSB_), INFβ (Eqn 1) and TREX1 (Eqn 2). First, particle transport Monte Carlo (FLUKA) simulations were performed for 6 MV x-rays, based on a phase space file from the CERR external beam dose calculation package initially modeled in VMC++,^23^ CNTS fast neutrons, as well as mono-energetic proton, helium-4 (^4^He), and carbon-12 (^12^C) particle beams with a range of 10 cm, all in water. The initial beam spot size was defined as 5 mm full width at half-maximum for all simulations, and with 0.2 mm voxel resolution. Next, user routines were developed within the FLUKA simulations to score the effective charge (z_eff_) and particle speed relative to the speed of light (β) within each voxel. Using an approach adapted from Stewart et al,^21^ the squared ratio of z_eff_ and β was applied to equation 3 to determine the RBE_DSB_, relative to colbalt-60 gamma rays.

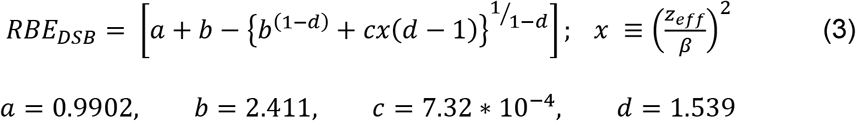

Lastly, depth-amplitude plots of the physical dose and RBE_DSB_ were applied to Eqns. 1 and 2 to determine and compare the variability in IFNβ and TREX1, as represented by RBE_IFNβ_ and RBE_TREX1_, across whole radiation fields.

## Results

### In vitro experiments

Increased levels of IFNβ were observed in MCC13 cells after x-ray and fast neutron irradiations. For each radiation type, a well-defined peak occurred in IFNβ production, which decreased at the highest doses assayed (Figure 1). Measurements show SARRP-irradiated cells exhibit an IFNβ peak at 14.0 Gy, while fast neutron-irradiated cells present a peak in IFNβ production at 5.7 Gy. The ratio of peak doses, defined here as RBE_IFNβ_, is 2.5 ± 0.2. The peak amplitudes of secreted IFNβ were not statistically different between the radiation modalities (P > 0.05). An upregulation of exonuclease TREX1 with radiation dose is shown in Figure 2. A linear increase of TREX1 with dose was measured, with neutron-irradiated cells expressing equivalent TREX1 at lower radiation dose. Fast neutron-irradiated cells upregulated TREX1 at a four-fold higher rate per unit dose than SARRP-irradiated cells (effective *RBE*_*TREX1*_ = 4.0 ± 0.1).

**Figure 1.**
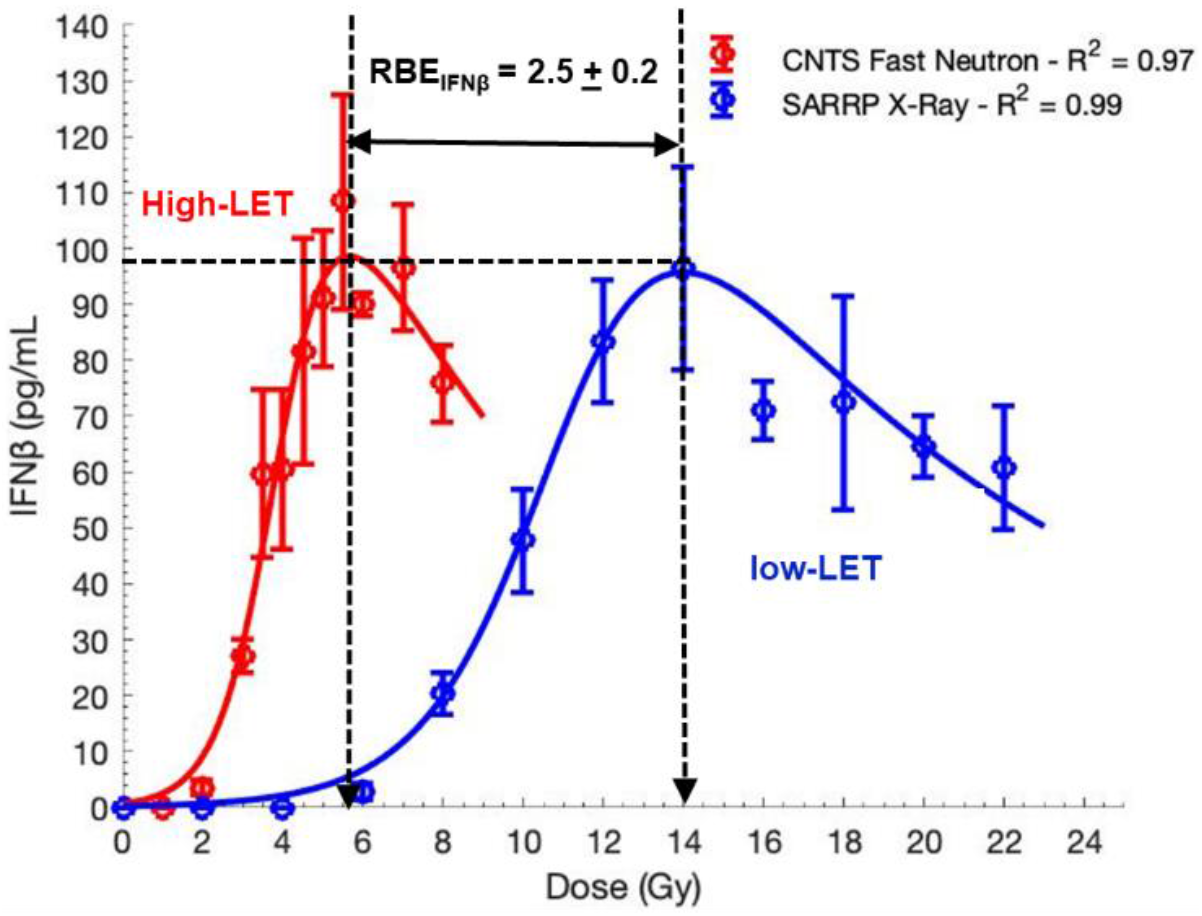
Secreted IFNβ dose response in MCC13 cells following SARRP x-ray irradiat ion and CNTS fast neutron irradiation by ELISA. The ratio of these peak doses is 2.5 ± 0.2, approximately equivalent to the RBE_DSB_ reported for CNTS neutrons^13^. There is no significant difference in the amplitude of peak IFNβ secretion (P > 0.05; paired t-test).

**Figure 2.**
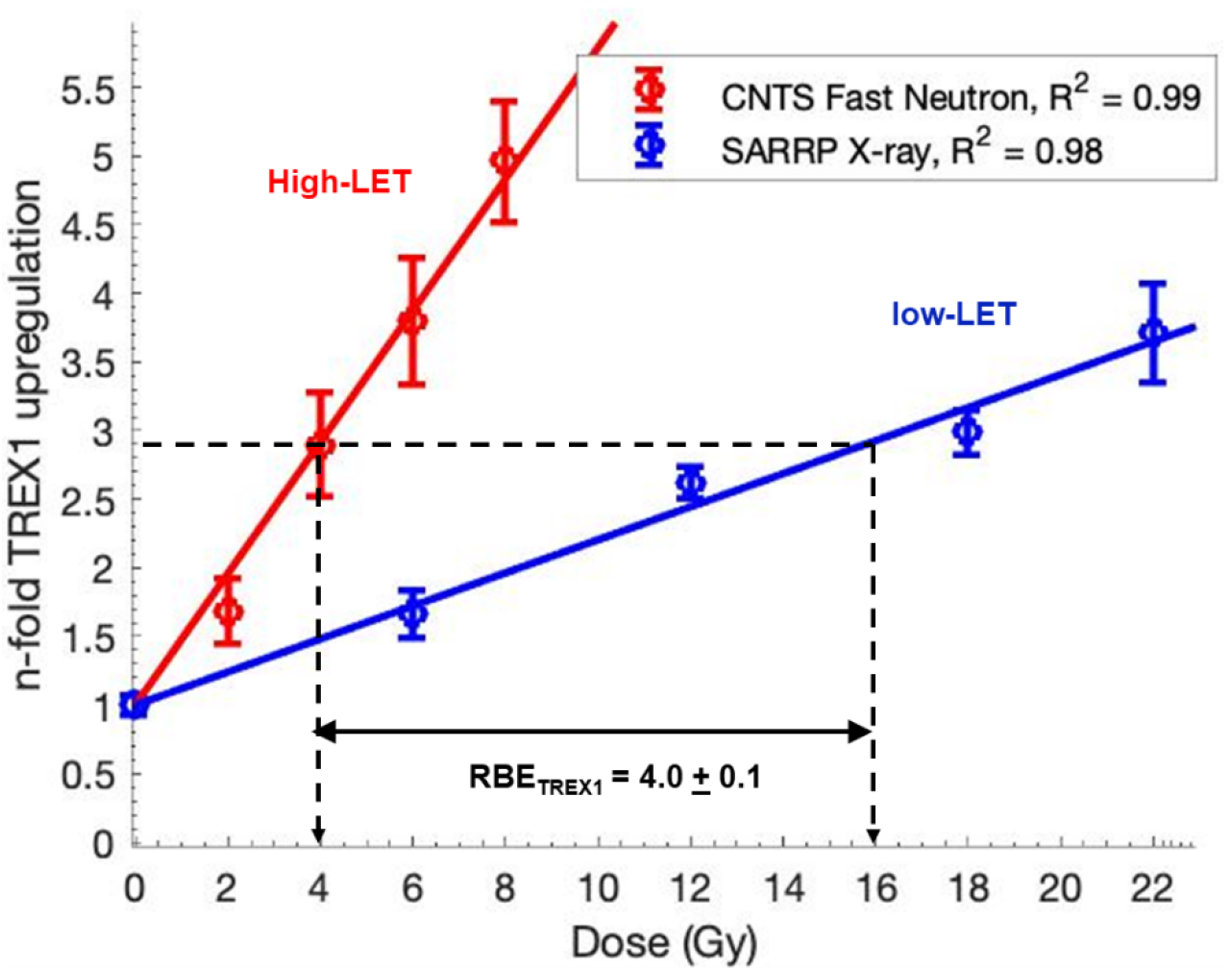
Measured TREX1 dose response in MCC13 cells following SARRP x-ray irradiation and CNTS fast neutron irradiation. Dose responses are normalized to background expression of TREX1. The ratio of slopes, i.e. the RBE_TREx1_, is 4.0 ± 0.1.

### Putative Relationship between DSB Induction and INFβ and TREX1 Response

Fit coefficients for IFNβ and TREX1 dose responses can be seen in Table 1, and RBE for a variety of low and high-LET radiation sources in Table 2. The R^2^ for all fits was > 0.97. Higher-LET radiation induces more DSB per unit dose than low-LET radiation, and our model predicts similar trends in IFNβ secretion and TREX1 upregulation (Figure 3). The ratio of doses that produce the same peak IFNβ response defines the potentially achievable RBE for IFNβ secretion, RBE_IFNβ_. We find that RBE_IFNβ_ is roughly equivalent to RBE_DSB_ obtained from Monte Carlo simulations, i.e., RBE_IFNβ_ ≅ RBE_DSB_ ≅ 2.3 to 2.5 (relative to SARRP x-rays). The effective RBE for TREX1 is about 2-fold larger than the RBE for IFNβ or DSB induction, RBE_TREX1_ ≅ 4.0 > RBE_IFNβ_ ≅ RBE_DSB_.

**Table 1.** Fit coefficients for radiation induced IFNβ and TREX1 for MCC13 cells. IFNβ measurements were fitted to a nonlinear function, and Trex was fitted to a first-order polynomial function using the MATLAB curve fitting toolbox (Eqn. 1 and 2).

|  | 220 kVp x-ray |  | CNTS Fast Neutron |  |
| --- | --- | --- | --- | --- |
| | IFNβ | Trex1 | IFNβ | Trex1 |
| **a** | 4.7E-03 | 0.10 | 7.2E-03 | 0.18 |
| **b** | 4.0E-06 | 1.00 | 2.4E-06 | 1.00 |
| **c** | 4.80 | -- | 1.50 | -- |
| **RBE<sub>DSB</sub>** | 1.17 |  | 2.70 |  |
| **R<sup>2</sup>** | 0.99 | 0.98 | 0.97 | 0.99 |

**Table 2.** Comparison of RBE_iFNβ_, predicted physical dose for optimal IFNβ, and FWHM of elevated IFNβ for a variety of radiation sources. Megavoltage photon and all charged particle values were determined using simulated data. RBE_DSB_ was modeled in MCDS relative to cobalt-60 for the default cell diameter and DNA content at normoxia. The IFNβ dose res ponse was modeled using Equation 1, which was applied to determine RBE_iFNβ_ relative to cobalt-60, the physical dose to produce optimal IFNβ. Note that due to continuous average energy loss across iontherapy beams, DNA damage coefficients, thus RBE_iFNβ_ are range dependent.

| Radiation Type | LET<br>(keV/αm) | RBE<sub>DSB</sub> | INFβ | TREX1 |
| --- | --- | --- | --- | --- |
|  |  |  | Peak Dose (Gy) | (n-fold upreg. / Gy) |
| Reference Radiation(s) |  |  |  |  |
| <sup>60</sup>Co γ-rays | 0.24 | 1.000 | 16.4 | 0.086 |
| 6 MV x-rays | 0.19 | 1.000 | 16.4 | 0.086 |
| SARRP x-rays | 3.87 | 1.169 | 14.0 | 0.117 |
| Proton* |  |  |  |  |
| Entry | 1.57 | 1.019 | 16.1 | 0.089 |
| Bragg Peak | 8.54 | 1.266 | 12.9 | 0.137 |
| Distal edge (50%) | 14.10 | 1.450 | 11.2 | 0.180 |
| <sup>4</sup>He<sup>2+</sup> ion* |  |  |  |  |
| Entry | 4.24 | 1.074 | 15.3 | 0.099 |
| Bragg Peak | 42.14 | 1.934 | 8.3 | 0.249 |
| Distal edge (50%) | 66.82 | 2.249 | 7.0 | 0.337 |
| <sup>12</sup>C<sup>6+</sup> ion* |  |  |  |  |
| Entry | 14.43 | 1.239 | 13.2 | 0.131 |
| Bragg Peak | 210.02 | 2.680 | 5.7 | 0.480 |
| Distal edge (50%) | 292.81 | 2.960 | 5.0 | 0.583 |
| CNTS Neutrons |  | 2.500 | 6.2 | 0.418 |
|  | 142.4** | 2.700 | 5.7 | 0.486 |
|  |  | 2.900 | 5.1 | 0.560 |

\* Monte Carlo simulation data for monoenergetic beams, 10 cm range in water
\*\* Effective CNTS neutron LET equivalent to a 140 MeV carbon ion (same value for RBEDSB)

**Figure 3.**
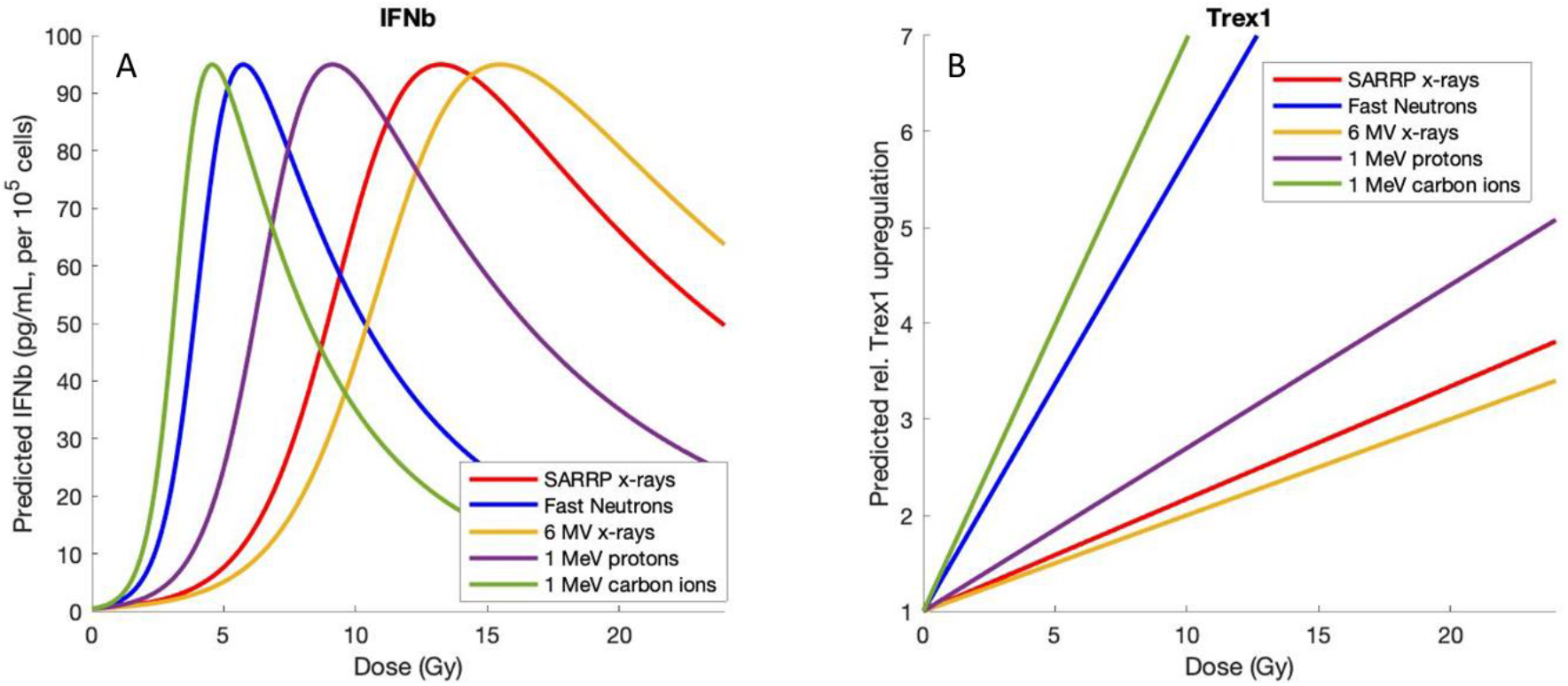
Modeled IFNβ and Trex1 regulation as a function of dose, for SARRP x-rays, fast neutrons, 6 MV x-rays, low-energy protons, and low-energy carbon ions. (A) IFNβ secretion as a function of dose for various radiation sources. High-LET radiation induces more DSB Gy^−1^ and have a larger RBE_DSB_, which parallels the increased efficiency of IFNβ secretion. (B) Trex1 upregulation as a function of dose for various radiation sources. Accumulation of DNA damage at lower doses enables Trex1 upregulation at lower doses. All fits to measured data use inputs for MCC13 cells (Table 1, 2).

### IFNβ secretion for representative clinical beams

RBE_IFNβ_ and physical dose values from FLUKA were applied to predict IFNβ secretion as a function of depth, as illustrated in Figure 4. The modeled level of IFNβ secretion is based on the Monte Carlo simulated value of the (absorbed dose) x RBE_DSB_ induction along pristine Bragg peaks and then scaled using Eq. (1) for the endpoint of IFNβ secretion. Uncharged radiation sources, such as low-LET 6 MV x-rays and fast neutrons, have a roughly constant RBE_DSB_ with depth as they propagate through tissue. For ion sources, however, the variability in LET and mean energy with depth drives an increase in RBE_DSB_ near the distal edge of their Bragg peaks resulting, we predict, in a more localized hotspot of IFNβ stimulation. Heavier ions have a sharper Bragg peak with respect to depth and a higher rate of IFNβ production at the Bragg peak relative to entrance levels (40-, 100- and 120-times for proton, ^4^He and ^12^C ion beams respectively), thus the doses for optimal pro-immunogenic response may become tighter with reduced activation proximally and distally, both biologically and dosimetrically.

**Figure 4.**
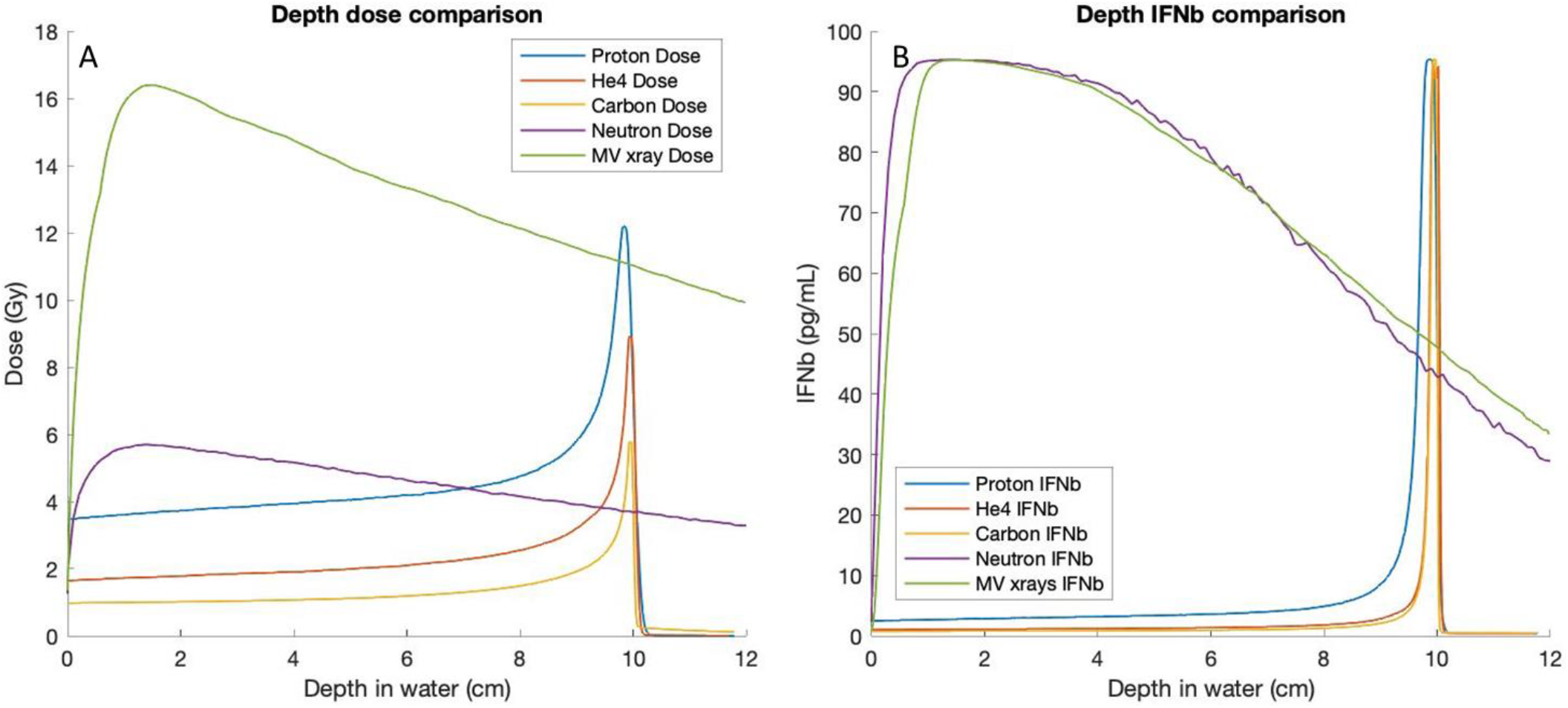
Monte Carlo model of IFNβ dose-response for 6 MV x-rays, CNTS fast neutrons, protons, alpha particles, and carbon ions. (A) Depth-dose plots for a variety of clinical beams, normalized to produce peak IFNβ in MCC13 cells treated at the depth of max dose. (B) IFNβ secretion as a function of depth in water for varying clinical beams. Delivered doses for ion sources can be normalized such that IFNβ is only stimulated around their Bragg peak; heavier ions can produce sharper regions of pro-immunogenic activity than indirectly ionizing x-rays or neutrons. Uncharged sources can stimulate elevated IFNβ more uniformly with depth.

## Discussion

While immunotherapy provides an exciting new treatment option, over half of solid tumors are unresponsive to immunotherapy alone.^24^ An increasingly advocated approach to enhance the efficacy of immunotherapy is to provide concurrent radiation to stimulate STING signaling and immunogenic cell death, which may be optimal only within a narrow radiation dose window.^25,26^ Limited prior data have explored the effects of radiation quality on radiation-enhanced immunogenicity. Here, we have demonstrated differential immunogenic responses, potentially via STING activation, in MCC13 cells by measuring the dose-response characteristics of IFNβ and TREX1 following irradiation by low-LET x-rays and high-LET fast neutrons, encapsulated in the computation of RBE_IFNβ_ and RBE_TREX1_. These data, in conjunction with Monte Carlo DNA damage modeling (MCDS and FLUKA simulations), were used to further investigate and generalize measured results for IFNβ and TREX1 regulation as a function of physical dose and RBE_DSB_. CNTS fast neutrons have a reported^21^ RBE_DSB_ of 2.3 + 0.2 relative to SARRP x-rays, which is equivalent to the RBE_DSB_ of 140 MeV ^12^C^6+^ ions (LET=142.4 keV/μm and a 0.57 mm range in water).^22^ The measured value in this study for RBE_IFNβ_ (2.5 ± 0.2) is remarkably close to the computed RBE_DSB_ value for CNTS neutrons relative to SARRP x-rays. This suggests that radiation-induced IFNβ secretion may scale with the number of initial DSB per cell. However, the RBE_TREX1_ value of 4.0 ± 0.1 was notably higher than RBE_IFNβ_ (and RBE_DSB_), indicating that TREX1 immunosuppression is more readily activated *in vitro* by higher-LET radiations than SARRP x-rays in MCC cells. Given the above observation, and that the peak magnitude of IFNβ production for x-rays and fast neutrons are statistically indistinguishable, our study may suggest more genomic DNA is entering the cytosol, per unit dose, of high-LET radiation when compared to the low-LET SARRP x-rays. The neutron RBE_IFNβ_ and RBE_TREX1_ reported here is higher than the published RBE for micronucleus (MN) induction of 2.0 ± 0.1, but smaller than the RBE for nucleoplasmic bridges of 5.8 ± 2.9.^27^ And, given that the mechanism of TREX1 upregulation in response to genotoxic stress remains poorly understood, further studies are needed to explain the putative mechanistic link relating DSB induction to the observed IFNβ and TREX1 responses.

The results from this study are consistent with several clinical case studies of MCC patients treated with fast neutrons and add additional insight into potential clinical improvements in radiation delivery to enhance radiation-induced immunogenicity. Schaub *et al.*^14^ report on the treatment of an 85-year-old man with chronic lymphocytic leukemia having progressive MCC with multiple tumors on the face despite prior low-LET radiation therapy and ongoing treatment with anti-PD1 pembrolizumab. The five most symptomatic lesions interfering with the patient’s vision were treated with two sequential neutron doses of 3 Gy separated by a few days while continuing pembrolizumab treatment. All five of the treated facial lesions demonstrated a complete response within 4 weeks. Remarkably, an additional 4 lesions located outside the treated areas also completely resolved, suggesting an adaptive anti-tumor immune response. Our models of MCC predict peak IFNβ production occurs near 17 Gy for 6 MV x-rays (Table 2) with little to no response at conventional x-ray doses (1.6-2 Gy) and most hypofractionated doses (6-8 Gy). With an RBE_IFNβ_ of 2.7+0.2 (relative to Co-60), peak IFNβ production for CNTS neutrons is anticipated to decrease to 5.7 Gy. At 3 Gy, IFNβ production is only 30% of its peak value. Our data suggest that a single dose of ~ 6 Gy of fast neutrons may be more effective at potentiating an out-of-field response in combination with ICI, with higher doses expected to provide little additional benefit and increase the potential for normal tissue complications. This is supported by prior studies that have observed abscopal effects in murine models of osteosarcoma following combined ICI and high-LET carbon ion therapy (5.3 Gy, a dose consistent with our models), however, direct comparison with low-LET radiation was not explored.^28^ Based on prior literature, repeat fractions are expected to further enhance IFNβ secretion, but the effects of fractionation^7^ are beyond the scope of this study.

Fast neutrons produced in the CNTS chiefly deposit dose through secondary low-energy (< 30 MeV on average) protons and (< 6 MeV) alpha particles, which offers a large RBE and immunogenic enhancement.^29^ One consideration of fast neutrons over charged particle therapy is that beyond the initial dose buildup region, their RBE/LET is roughly uniform with depth. Less variability across a beam allows more simple treatment planning and optimization purely based on physical dose. However, for ion therapy beams, the narrow spatial regions of high-LET radiation at the tip of a Bragg peak may be used advantageously to more-precisely sculpt regions where pro-immunogenic effects are desired. Although preliminary, our models enable simulation of such effects across entire radiation fields, which may be useful for guidance of future studies and clinical trials, if supported by further *in vivo* validation of these effects. Tables 1 and 2 provide a general framework for calculating RBE_IFNβ_ and RBE_TREX1_ for x-rays and fast neutrons, as well as for common ions superficially and at their end of range. The results in Figure 4 illustrate how an IFNβ-stimulatory radiation field can be constructed. The RBE_IFNβ_ factor can be used to define the dose that will stimulate peak IFNβ at depth for a specific radiation field (MV, proton, alpha, carbon ion). We observed peak-to-entrance IFNβ ratios to be approximately 40 to 120 for charged particle sources (Figure 4), a factor of ten higher compared to physical dose peak-to-entrance. Due to this nonlinear response (Figure 1), the off-target dose and IFNβ production can be kept significantly lower.

A potential factor suppressing IFNβ is the increase in TREX1 with dose. Measurements from this study show a faster rise in TREX1 with increasing dose for high-LET neutrons than for low-LET x-rays (Figure 2). This rapid increase in TREX1 with increasing dose and trends in RBE (RBE_TREX1_ > RBE_IFNβ_ ≅ RBE_DSB_), as seen in Figures 1 and 2, collectively imply a potential dampening effect of TREX1 on IFNβ that is larger for high LET neutrons than for low LET x-rays. One potential explanation for this observation is that DNA fragment length may play a role in regulating immune signaling. Prior works have demonstrated a dependence on both DNA strand length and concentration on cytosolic DNA sensing for immunogenic signaling through cGAS-STING^30^ and, on average, high LET radiations tend to produce smaller DNA fragments than those produced by low LET radiations.^31,32^ The data in this study suggest small cytosolic DNA fragments preferentially produced by high LET radiations may be more efficiently detected and degraded by TREX1 than large DNA fragments produced by low LET radiations. Regardless of the underlying mechanisms, our findings suggest an enhanced immunogenic response may be further realized at lower doses through TREX1 targeted interventions.^33^

One limitation of this study is its breadth. Our experiments focused on immune responses in one Merkel carcinoma cell line (MCC13). Some care in evaluating the histopathological characteristics of other cell lines should be considered, largely due to the rarity of cultured cell lines having intact STING signaling. For example, deficiencies in cGAS, a cytosolic double-stranded DNA sensor, are common in colon adenocarcinoma cell lines, which frequently occur following hypermethylation of promoter regions, and can in part be restored using demethylation agents.^34^ The investigation of additional tumor cell lines with intact cGAS-STING will shed light on whether or not the observations described here for MCC13 cells are more generalizable.

The effects of varying dose rates on radiation-induced STING signaling is a planned further study, particularly in the context of FLASH ultra-high dose rate radiation therapy. *In vivo* study of radiation delivered at ultra-high dose rates (> 40 Gy/s) has been demonstrated to preferentially spare normal tissues while preserving tumor control at equivalent physical doses, relative to conventional dose rates.^35^ A common theory for this differential effect is that nearly instantaneous radiation delivery damages fewer circulating lymphocytes, limiting systemic immunosuppression. This has, in part, been confirmed *in vivo*, where enhanced CD8+ T cell accumulation has been observed following irradiation of immunocompetent mouse tumor models.^36^ However, more studies are needed to further explore the inflammatory response of cancer cells following FLASH radiation.

To conclude, we have presented a comparative analysis of pro-immunogenic signaling initiated within MCC13 cancer cells and its suppression from high-LET neutrons relative to low LET x-rays. The findings from this study provide three important results. First, high-LET radiation produces peak levels of IFNβ at lower doses compared to low-LET radiation (Figure 1), consistent with Monte Carlo models of DSB induction. Second, the rapid increase in TREX1 with dose and trends in RBE (RBE_TREX1_>RBE_IFNβ_ ≅ RBE_DSB_) indicate fast neutrons have a larger dampening effect on IFNβ than x-rays, and TREX1 targeted interventions may further the immunogenic response and reduce treatment doses.^33^ Third, the LET from the CNTS is consistent with that found in the Bragg peak of carbon ion beams and, therefore, our results may provide insights into the usefulness of carbon ions and other high LET radiations for the stimulation of immunogenic responses. Results from this study are useful for the validation of mathematic models to predict depth-IFNβ-curves for low and high-LET radiations and help guide optimization IFNβ activation in tumor targets relative to healthy tissue. Further work is necessary to test the variability in radiation-induced STING signaling across cell lines *in vitro*, although this work provides useful insight and data to understand and translate in vivo studies of IFNβ and TREX1 to the activation of immune responses *in vivo*.

## Acknowledgements

The authors would like to thank Robert Emery, David Argento, Eric Dorman, and Marissa Kranz of the University of Washington Medical Cyclotron Facility and Vadim Moskvin of the Radiation Oncology Department at St. Jude Children’s Research Hospital for their support of the reported studies.

